# Granulocyte-colony stimulating factor (G-CSF) enhances cocaine effects in the nucleus accumbens via a dopamine release-based mechanism

**DOI:** 10.1101/2021.07.19.452968

**Authors:** Lillian J. Brady, Kirsty R. Erickson, Kelsey E. Lucerne, Aya Osman, Drew D. Kiraly, Erin S. Calipari

**Affiliations:** Department of Pharmacology, Vanderbilt University, Nashville, TN 37232, USA; Vanderbilt Center for Addiction Research, Vanderbilt University, Nashville, TN 37232, USA; Vanderbilt Brain Institute, Vanderbilt University, Nashville, TN 37232, USA; Nash Family Department of Neuroscience, Icahn School of Medicine at Mount Sinai, New York, NY 10029, USA; Friedman Brain Institute, Icahn School of Medicine at Mount Sinai, New York, NY 10029, USA; Department of Psychiatry, Icahn School of Medicine at Mount Sinai, New York, NY 10029, USA; Seaver Autism Center for Research and Treatment, Icahn School of Medicine at Mount Sinai, New York, NY 10029, USA; Vanderbilt Institute for Infection Immunology and Inflammation, Vanderbilt University School of Medicine, Nashville, TN 37232, USA; Department of Molecular Physiology and Biophysics, Vanderbilt University, Nashville, TN 37232, USA; Department of Psychiatry and Behavioral Sciences, Vanderbilt University School of Medicine, Nashville, TN 37232, USA

**Author notes:** **Corresponding Authors Erin S. Calipari, PhD**, Assistant Professor, Department of Pharmacology, Department for Molecular Physiology and Biophysics, Department of Psychiatry and Behavioral Sciences, Vanderbilt Center for Addiction Research, Vanderbilt Brain Institute, Vanderbilt University School of Medicine, 865F Light Hall, 2215 Garland Avenue, Nashville, TN 37232, **Drew D. Kiraly, MD, PhD**, Assistant Professor, Department of Psychiatry, Nash Family Department of Neuroscience, Friedman Brain Institute, Icahn School of Medicine at Mount Sinai, 1 Gustave L Levy Pl - Box 1230, New York, NY 10029. Denotes equal contribution (Lillian J. Brady, Kirsty R. Erickson). Denotes co-corresponding and equal contribution (Drew D. Kiraly, Erin S. Calipari).

## Abstract

Cocaine use disorder is associated with alterations in immune function including altered expression of multiple peripheral cytokines in humans - several of which correlate with drug use. Individuals suffering from cocaine use disorder show altered immune system responses to drug-associated cues, highlighting the interaction between the brain and immune system as a critical factor in the development and expression of cocaine use disorder. We have previously demonstrated in animal models that cocaine use upregulates expression of granulocyte colony stimulating factor (G-CSF) - a pleiotropic cytokine - in the serum and the nucleus accumbens (NAc). G-CSF signaling has been causally linked to behavioral responses to cocaine across multiple behavioral domains. The goal of this study was to define whether increases in G-CSF alter the pharmacodynamic effects of cocaine on the dopamine system and whether this occurs via direct mechanisms within local NAc microcircuits. We find that systemic G-CSF injection increases cocaine effects on dopamine terminals. The enhanced dopamine levels in the presence of cocaine occur through a release-based mechanism, rather than through effects on the dopamine transporter - as uptake rates were unchanged following G-CSF treatment. Critically, this effect could be recapitulated by acute bath application of G-CSF to dopamine terminals, an effect that was occluded by prior G-CSF treatment, suggesting a similar mechanistic basis for direct and systemic exposures. This work highlights the critical interaction between the immune system and psychostimulant effects that can alter drug responses and may play a role in vulnerability to cocaine use disorder.

## Introduction

There has been growing interest in recent years in the interaction between immune systems and neuronal function, and emerging evidence suggests that immune dysfunction plays a critical role in the etiology of neuropsychiatric disease, including substance use disorders^1–3^. Accordingly, immune dysregulation has been implicated in addiction, and only recently have studies begun to examine the mechanistic link between altered immune function and the pathology underlying addictive disorders^2,4–8^. Granulocyte-colony stimulating factor (G-CSF), a cytokine, is regulated by cocaine in rodents^6^ and humans^9^, and is a potent modulator of behavioral plasticity induced by repeated cocaine exposure where it increases cocaine self-administration under low effort schedules of reinforcement, enhances motivation for cocaine, increases cocaine conditioned place preference, increases sensitization, and alters drug seeking in both males and females^5,6,10^. While the behavioral effects are well-characterized, the neural basis for how this cytokine enhances stimulant effects remains to be elucidated.

At the hub of dysregulation of reward and motivation in cocaine use disorder is the mesolimbic dopamine system. Dopamine projections from the ventral tegmental area (VTA) to the nucleus accumbens (NAc) are critical for cocaine reinforcement and the development of cocaine use disorder^11–14^. There is evidence that proinflammatory cytokines can alter mesolimbic dopamine function by regulating dopamine clearance, dopamine receptor expression, and/or altering synthesis and releasable pool content^7,15^. This suggests that cytokines may be acting through multiple synaptic mechanisms, especially via the dopamine system, to exert their effect on reward-related behaviors. Interestingly, G-CSF alters basal evoked dopamine release at terminals in the NAc^4^, and previous work in females, suggests that G-CSF can alter selective pharmacodynamic properties of cocaine, however, this was only under certain conditions^5^. The goal of this study was to understand whether G-CSF enhances the effects of cocaine in males and define the mechanistic basis by which this occurs.

Stimulant drugs of abuse, such as cocaine, increase dopamine levels in the NAc via direct actions at dopamine terminals. First, cocaine reduces the rate of dopamine clearance via dopamine transporter (DAT) blockade, thus increasing the relative levels and duration of synaptic dopamine signaling^16–18^. Additionally, cocaine has been shown to enhance release from terminals through exocytotic mechanisms that depend on synapsin^19^. Thus, cocaine acts both on release and clearance mechanisms at the terminal to exert its global effects on tonic dopamine levels in reward-related brain regions. Because release and clearance processes have been shown previously to be regulated by a variety of peripheral factors in the absence of cocaine^5,20^, cocaine effects at terminals are well situated to be modulated and regulated via similar mechanisms. Currently, it is not clear if effects of G-CSF on the mesolimbic dopamine system are selectively through release-based mechanisms - as have been observed at baseline - or through alterations in transporter function that enhance cocaine affinity for the dopamine transporter.

Using fast scan cyclic voltammetry to record subsecond dopamine release from terminals in the NAc, we investigated how systemic G-CSF treatment or direct bath application to terminals alters the pharmacodynamic properties of cocaine. We find that G-CSF enhances dopamine release in the presence of cocaine. This effect occurs through a release-based mechanism, rather than through effects on the dopamine transporter. Importantly, this release-based mechanism could not be explained purely by changes to evoked release at baseline and represented additive enhancement of cocaine effects on release. Critically, this effect could be recapitulated by acute bath application of G-CSF to dopamine terminals, highlighting direct microcircuit regulation in the NAc is sufficient for this to occur. Together, this work highlights the critical interaction between the immune system and psychostimulant effects that can alter drug responses and likely plays a role in vulnerability to cocaine use disorder.

## Materials and Methods

### Subjects

Male C57BL/6 J mice (7wks old ∼20–25 g; Jackson Laboratories, Bar Harbor, ME; SN: 000664) were housed in the animal facilities at Vanderbilt University/Vanderbilt University Medical Center. Mice were maintained on a 12:12 h reverse light/dark cycle (0700 hours lights off; 1900 hours lights on) and were housed three to five per cage. Animals were provided with food and water ad libitum throughout the experiments. All animals were maintained according to the National Institutes of Health guidelines in Association for Assessment and Accreditation of Laboratory Animal Care accredited facilities. All experimental protocols were approved by the Institutional Animal Care and Use Committee at Vanderbilt University/Vanderbilt University Medical Center. Experimenters were blind to experimental groups and order of testing was counterbalanced between groups.

### Drugs

For slice experiments, cocaine (NIDA Drug Supply Program) or granulocyte colony stimulating factor (G-CSF; GenScript, Piscataway, NJ) was dissolved in artificial cerebrospinal fluid on the day of the experiment and applied to brain slices via bath perfusion. For injections, granulocyte colony stimulating factor (G-CSF; GenScript, Piscataway, NJ) was dissolved in 0.9% sterile saline and administered intraperitoneally (IP).

### G-CSF Treatment

Animals were treated with G-CSF as described previously^4–6,10^. Briefly, G-CSF was injected via an IP injection twice: 24 hours before voltammetric recordings and again one hour before recordings. This dose and paradigm was chosen as it is consistent with our previous work, and results in comparable systemic levels to those induced by chronic cocaine exposure^4–6,10^.

### Fast Scan Cyclic Voltammetry

*Ex vivo* fast-scan cyclic voltammetry (FSCV) was used to characterize dopamine release in the NAc and its response to cocaine as described previously^20^. A vibrating tissue slicer was used to prepare 300 μM-thick coronal brain sections containing the NAc, which were immersed in oxygenated aCSF containing (in mM): NaCl (126), KCl (2.5), NaH2PO4 (1.2), CaCl2 (2.4), MgCl2 (1.2), NaHCO3 (25), glucose (11), L-ascorbic acid (0.4) and pH was adjusted to 7.4. The slice was transferred to the testing chambers containing aCSF at 32 °C with a 1 ml min^-1^ flow rate. A carbon fiber microelectrode (100– 200 μM length, 7 μM radius) and bipolar stimulating electrode were placed into the NAc core in close proximity to one another (**Fig 1B**). Dopamine release was evoked by a single electrical pulse (350 μA, 4 ms, monophasic) applied to the tissue every 5 min. Extracellular dopamine concentration was recorded by applying a triangular waveform (−0.4 to +1.3, to -0.4 V versus Ag/AgCl, 400 V s^-1^) to the carbon fiber recording electrode. Once the peak of evoked dopamine release was stabilized (three collections with <10% variability), cocaine experiments commenced. Recording electrodes were calibrated by recording responses (in electrical current; nA) to a known concentration of dopamine (3 μM) using a flow-injection system. This was used to convert an electrical current to a dopamine concentration.

**Figure 1.**
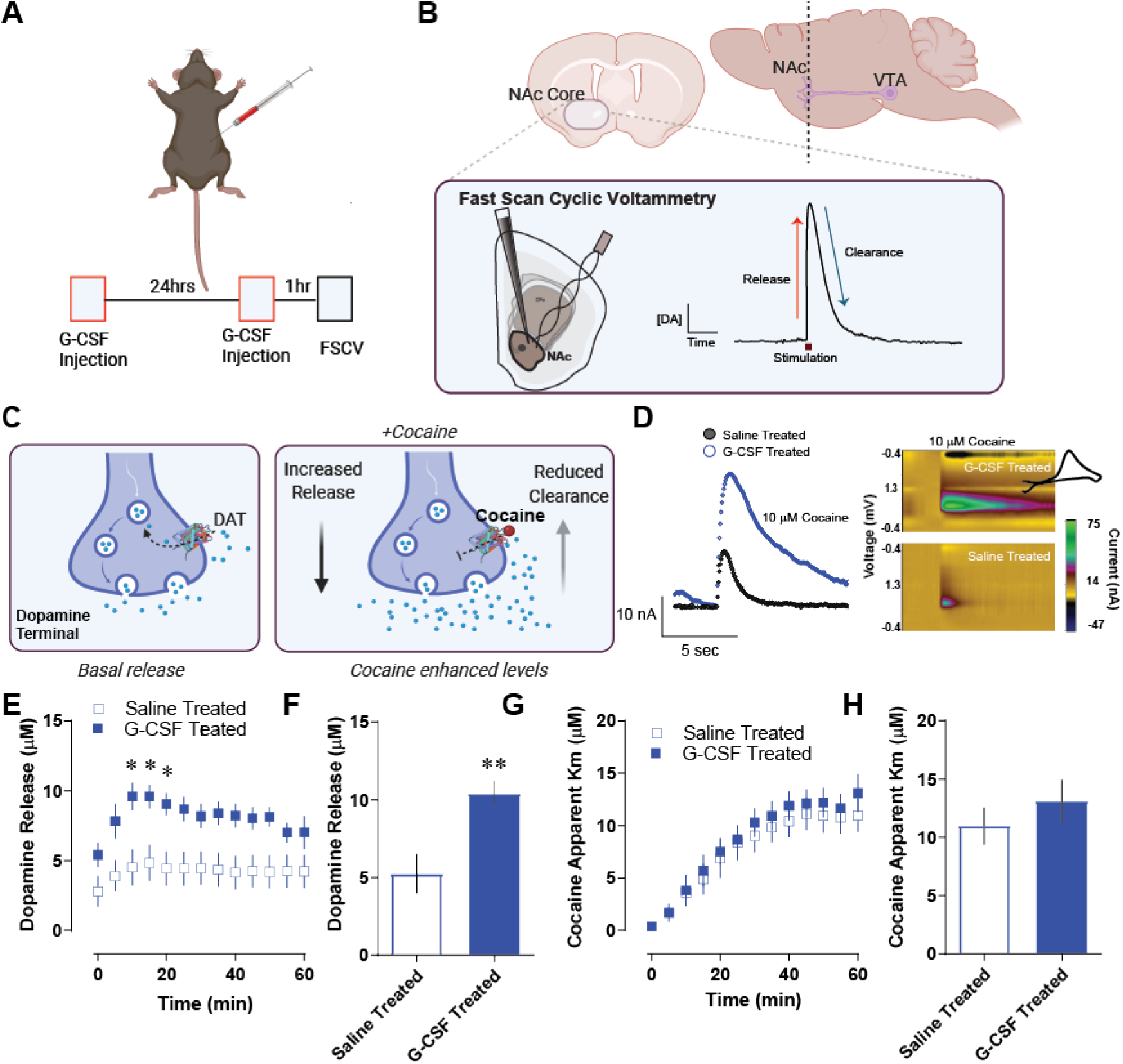
Granulocyte colony stimulating factor enhances cocaine-induced dopamine release in the nucleus accumbens core. **A**. Male C57BL6/J mice were injected with G-CSF (50μg/kg, IP) twice – 24 hours before recording and again one hour before recording. **B**. Fast scan cyclic voltammetry was done in the NAc core. Dopamine release was evoked from dopamine terminals via an electrical stimulation (n = 6 per group). Dopamine is recorded at 10 Hz via a carbon fiber microelectrode, which allows for sub second dopamine monitoring (inset, right) and the dissociation of dopamine release, from uptake-related measures. **C**. Cocaine increases extrasynaptic dopamine levels within the NAc via multiple mechanisms. Its canonical effects occur via inhibiting the dopamine transporter (DAT) and inhibiting reuptake; however, it also acts to promote exocytotic release^19^. These studies were designed to specifically parse release-based mechanisms from clearance-based and DAT-mediated mechanisms. **D**. Pre-treatment with G-CSF before recording sessions resulted in enhanced effects of cocaine on dopamine terminals in the NAc. *Left*, Current versus time plots showing measured dopamine levels during the application of 10µM cocaine to the slice. *Right*, color plots showing dopamine (green) at its oxidation potential as well as a cyclic voltammogram (*inset*) showing its characteristic redox signature of dopamine. **E**. Group data showing peak dopamine release over time after the bath application of 10µM cocaine in saline or G-CSF treated mice. **F**. The maximum evoked release in the presence of cocaine in each subject. Extrasynaptic dopamine levels in the presence of cocaine were enhanced following systemic G-CSF treatment. **G**. Michaelis-Menten modeling was used to assess the apparent km of cocaine for the dopamine transporter (a measure of relative affinity of cocaine for DAT). This was plotted over time following cocaine bath application. **F**. The apparent Km from each animal was plotted following stabilization. G-CSF did not affect cocaine effects at the dopamine transporter, suggesting that its effects are specific to release. Data represented as mean +/- S.E.M., ^*^ *p* < 0.5, ^**^ *p* < 0.01.

To assess the effects of cocaine on dopamine terminals in the NAc core, cocaine was bath-applied at a concentration of 10µM until the effects on clearance and release stabilized (45 minutes to 60 minutes). Each collection over the ∼60minute period was analyzed to assess area under the curve (AUC), Apparent Km, Tau, and peak height as described in Yorgason et al., 2012.

### Voltammetric data analysis

Demon voltammetry and analysis software was used for all analysis of FSCV data^21^. Data were modelled either using Michaelis–Menten kinetics to determine dopamine release and apparent Km or via analyzing peak and decay kinetics - for AUC and Tau.

For baseline, peak and return to baseline measurements computations were based on user-defined positions on current traces. Tau values were determined from exponential fit curves based on the peak and the post-peak baseline using a least squares constrained exponential fit algorithm (National Instruments). Peak height, (in nA), and [DA] (in μM) were determined by the peak of the evoked signal, while AUC was calculated as the numeric integration of the area between the data point immediately preceding the stimulations and the point at which the signal returned to baseline, using a trapezoidal rule.

For Michaelis-Menten modeling, we assumed that all parameters were floating and found the best fit line for each data point as described in Calipari et al. (2017)^20^. The Michaelis-Menten based parameters, [DAp], Vmax, and Km, were used to evaluate release and uptake kinetics using the following equation^22–24^:

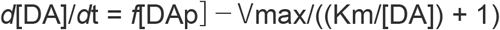

Where [DA] is the instantaneous extracellular concentration of dopamine released, *f* is the stimulation frequency, [DAp] is the release rate constant (expressed as concentration of dopamine released per stimulus pulse) and Vmax and Km are Michaelis-Menten clearance rate constants. To assess cocaine effects on the dopamine terminal, a baseline Vmax was determined individually for each subject. Following cocaine bath perfusion, Vmax was held constant for the remainder of the experiment and cocaine-induced changes in uptake were attributed to changes in apparent Km - the effect of cocaine on DAT as inferred by the change in dopamine clearance relative to baseline.

## Statistical Analysis

All statistical analysis was performed using GraphPad Prism. Pairwise comparisons were performed using a two-tailed Student’s *t*-test with Welch’s correction when appropriate. One factor comparisons were performed using one-way ANOVA and Sidak’s post-hoc tests. 2 × 2 comparisons were performed using two-way ANOVA with repeated measures and Holm-Sidak’s post-hoc tests. Correlational analyses were performed using Pearson’s correlation analysis. Alpha values for all analyses were set to 0.05.

## Results

### G-CSF increases cocaine effects on dopamine release in the nucleus accumbens

First, we aimed to define how increasing G-CSF levels alters the pharmacodynamic effects of cocaine on the dopamine system in the NAc core. Male C57BL/6 J mice were injected IP with G-CSF (50μg/kg) as outlined in **Figure 1A**. This injection paradigm results in G-CSF levels that are comparable to the increases we have seen previously following repeated cocaine exposure^6^. Subsequently coronal slices were prepared, and fast scan cyclic voltammetry was run to determine subsecond dopamine release and clearance dynamics in the NAc core (**Fig 1B**). Voltammetry is a powerful rapid-sampling technique that allows for the dissociation of release from clearance-based mechanisms following evoked signals (**Fig 1B, inset**). This is particularly important for these studies as stimulants, including cocaine, have been shown to exert their effects through modulating both release and DAT-mediated clearance mechanisms (**Fig 1C**)^16,17,19^.

We found that G-CSF treatment increased cocaine effects on dopamine signals in the NAc core (**Fig 1D**). When we analyzed release effects, we found a significant overall effect of G-CSF treatment of cocaine [**Fig 1E**, Repeated measures two-way ANOVA, significant effect of G-CSF treatment, F (1, 10) = 7.94, *p* < 0.05]. Indeed, G-CSF injection increased the maximum amount of dopamine released in the presence of cocaine [**Fig 1G**, Student’s two-tailed t-test, t (10) = 3.44, *p* < 0.01]. While the effects on release were robust, there was not an effect on any clearance-based measures, (**Fig 1G-H**). Apparent Km is a measurement of the affinity of cocaine for DAT and there was no significant effect at any time-point following G-CSF treatment. Together, G-CSF increased cocaine effects on extracellular dopamine levels in the NAc which were selective to effects on peak evoked dopamine release, and were not mediated by effects at DAT.

### Cocaine effects on dopamine levels are correlated with release-based measurements, and not clearance

Voltammetric recordings allow for a complex analysis to dissociate release and clearance-based regulation of dopaminergic transmission (**Fig 2A**). In these experiments, dopamine was evoked by an electrical pulse applied to the brain slice. This results in rapid release followed by a clearance phase, which has been shown previously to be dependent on DAT (**Fig 2A**). Release and clearance together combine to determine the timing, magnitude, and duration of the dopamine response. Therefore, we analyzed the total area under the curve to represent the entire dopaminergic response (**Fig 2B**). We also parsed release and clearance-based measurements by plotting the peak of dopamine release and Tau [the time in seconds it takes to return to ⅔ of peak height (**Fig 2C**)].

**Figure 2.**
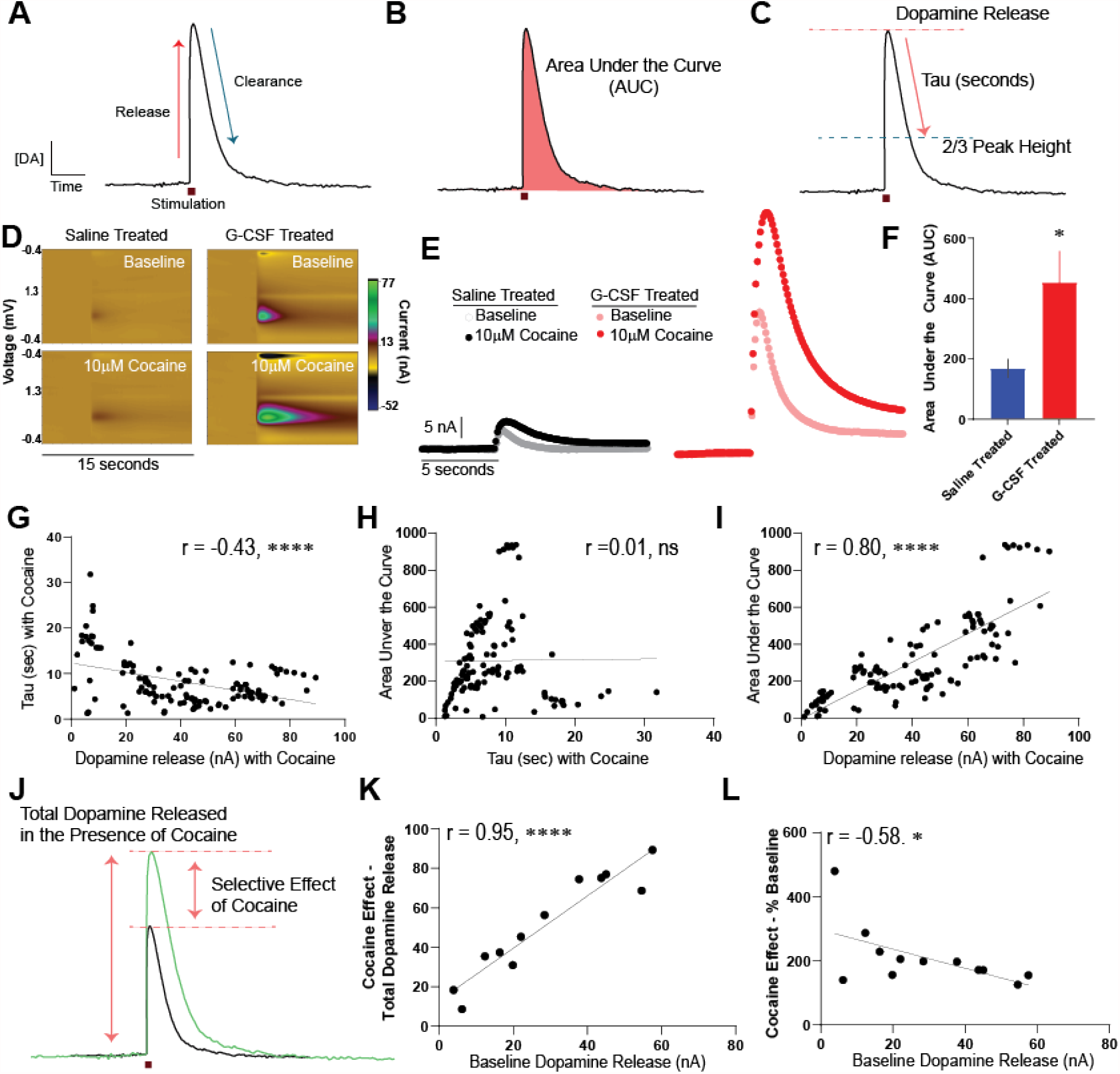
G-CSF enhancement of cocaine effects on the dopamine system is via a release-based mechanism. **A**. Example of an evoked dopamine signal over time. Significant work has defined what biological factors dictate specific components of the curve, with the rising phase being mediated by active dopamine release and the falling phase via DAT-mediated clearance. Changes in either factor can contribute to changes in the total timing and amount of dopamine in the extrasynaptic space. **B**. The magnitude of dopamine response, which is a function of both release and clearance mechanisms, can be assessed via area under the curve measurements (AUC) which represents the total effect of cocaine on the dopamine signal. **C**. Release and clearance-based measurements were parsed by assessing the total dopamine release (determined as the peak signal), and tau (the time in seconds it takes for the signal to go from peak to 2/3 of peak height). These analyses and comparisons were done for all groups. **D**. Representative color plots showing evoked dopamine responses at baseline (top) and following bath application of 10µM cocaine (bottom) following saline or G-CSF injection. **E**. Current versus time plots showing all groups at baseline and following bath application of cocaine on the same scale. *Left*, saline-injected animals before and after cocaine. *Right*, G-CSF injected animals. **F**. AUC analysis of the cocaine curves for saline treated or G-CSF treated animals showing that G-CSF treatment enhances cocaine effects. **G**. A series of correlations were run in order to parse the factors contributing to these dopamine signals. Correlation of Tau in the presence of cocaine and release in the presence of cocaine, showing that they are weakly negatively correlated. This means that more time to return to baseline is associated with less release. Data represented as mean +/- S.E.M., ^*^ *p* < 0.5, ^****^ *p* < 0.0001.

Cocaine increased dopamine responses in both saline animals and animals that had been treated with G-CSF (**Figure 2D, E)**. The AUC in the presence of 10µM cocaine was increased by prior G-CSF treatment relative to saline treated animals [**Fig 2F**, two-tailed Student’s t-test, t (9) = 2.42, *p* < 0.05]. Next, we conducted a series of correlations to determine the contributors to the variance in signal.

First, we correlated Tau (a measure of the clearance rate) in the presence of cocaine with the peak dopamine release in the presence of cocaine (**Fig 2G**). Increased Tau corresponds to an increase in the effect of cocaine on DAT inhibition. There was a negative correlation between the peak dopamine release in the presence of cocaine and cocaine effects on clearance (Pearson’s r = -0.44, R^2^ = 0.19, *p* < 0.0001). This demonstrates that the increased effects on release cannot be explained by cocaine effects at the DAT as the directionality is opposite from what would be expected if the cocaine effects on release could be attributed to cocaine’s canonical mechanism of action at DAT. Next, we correlated the AUC in the presence of cocaine - which accounts for the total magnitude of any clearance and release effects together - with Tau to determine if the AUC effects could be explained by differences in effects on clearance-based mechanisms (**Fig 2H**). There was no relationship between AUC and Tau (Pearson’s r = 0.01, R^2^ = 0.00009, not significant), indicating that the effects of G-CSF on overall dopamine responses cannot be explained by effects on clearance/DAT. Lastly, we correlated the AUC in the presence of cocaine with the peak of dopamine release in the presence of cocaine, which was significantly correlated (**Fig 2I**, Pearson’s r = 0.80, R^2^ = 0.63, *p* < 0.0001).

Finally, to understand the relationship between baseline release and cocaine effects on release we analyzed cocaine effects on release in two ways (**Fig 2J**). The first was taking the overall dopamine release in the presence of cocaine, which reflects the overall dopamine present in the NAc core. The second, was looking at the relationship between baseline release and the percent increase in release in the presence of cocaine. This second form of analysis looks at the overall cocaine effect and separates it from global changes in release across animals. First, we found that the cocaine effect on release when measured in terms of total peak dopamine signal (reported in current, nA) was significantly correlated with the amount of dopamine at baseline within each animal (reported in current, nA) (Pearson’s r = 0.95, R^2^ = 0.90, *p* < 0.0001). However, the effects of cocaine when normalized to release were weakly inversely correlated across the animals in the study (Pearson’s r = -0.58, R^2^ = 0.87, *p* < 0.05); thus, increases in dopamine release that are mediated by G-CSF injections at baseline^4^, cannot explain the enhanced effects of cocaine that were observed within this study.

### Acute bath application of G-CSF enhances cocaine effects on release

While the previous series of experiments showed that G-CSF enhances the effects of cocaine on dopamine terminals, we next wanted to understand if these effects are through direct actions within the NAc. To this end, we bath-applied 10nM G-CSF or vehicle to brain slices for 60 minutes before the subsequent application of 10µM cocaine (**Fig 3A,B**).

**Figure 3.**
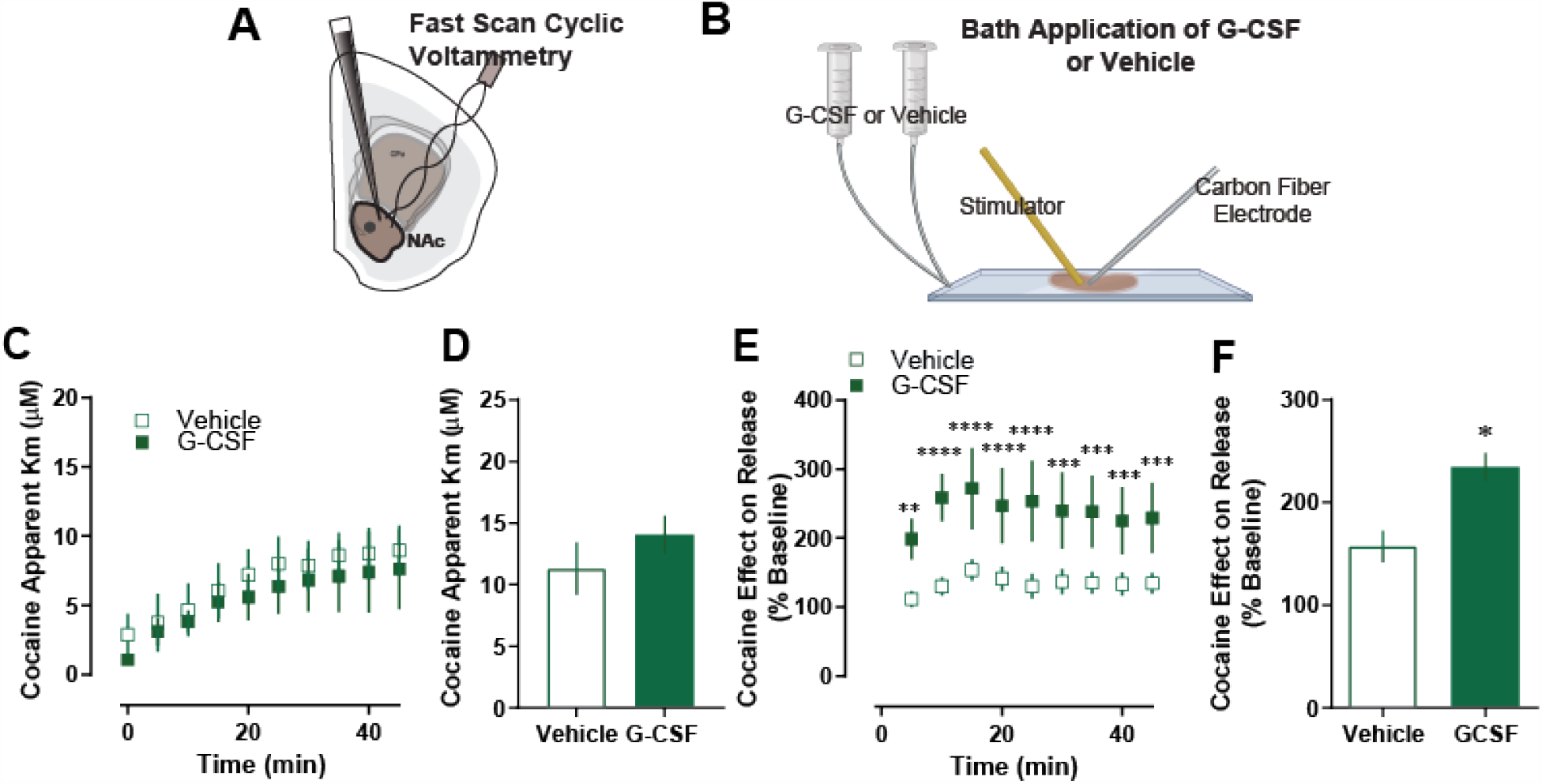
G-CSF enhances cocaine effects through direct actions in the nucleus accumbens. **A**. In these control animals, fast scan cyclic voltammetry was run. **B**. G-CSF or vehicle was applied via bath application to the NAc slices for one hour prior to cocaine application (n = 5 per group). **C**. G-CSF bath application had no effect on cocaine effects at DAT as measured by apparent Km. **D**. Group averages also showed no difference in apparent Km measures. **E**. Cocaine effect on dopamine release as presented as a percent of baseline. **F**. G-CSF bath application increased cocaine effects on dopamine release. Data represented as mean +/- S.E.M. ^*^, *p* < 0.05.

Direct application of 10nM G-CSF to dopamine terminals replicated the effects of systemic G-CSF injection. Acute G-CSF bath application did not have any effect on dopamine release or clearance at baseline, as we have reported previously. It also had no effect on cocaine effects at the dopamine transporter (**Fig 3C, D**), but did increase the effect of cocaine on evoked dopamine release over time [**Fig 3E**, two-way repeated measures ANOVA; F (1, 4) = 9.36, *p* < 0.05] and in relative magnitude [**Fig 3F;** Students t-test, t (4) = 3.87, *p* < 0.05), suggesting that the effects of G-CSF on cocaine effects occur directly in the NAc.

### Prior systemic G-CSF injection occludes the effects of G-CSF bath application on terminal effects

Finally, we tested whether G-CSF bath application had an effect following G-CSF injection. To this end, we injected animals with G-CSF (**Fig 4A**) before preparing slices for voltammetry. Subsequently, we bath-applied 10nM G-CSF or vehicle to brain slices for 60 minutes before the application of 10µM cocaine (**Fig 4B**). Acute G-CSF bath application did not have any effect on cocaine effects at the dopamine transporter (**Fig 4C, D**), or on evoked dopamine release (**Fig 4E, F**) in animals that were previously treated with G-CSF.

**Figure 4.**
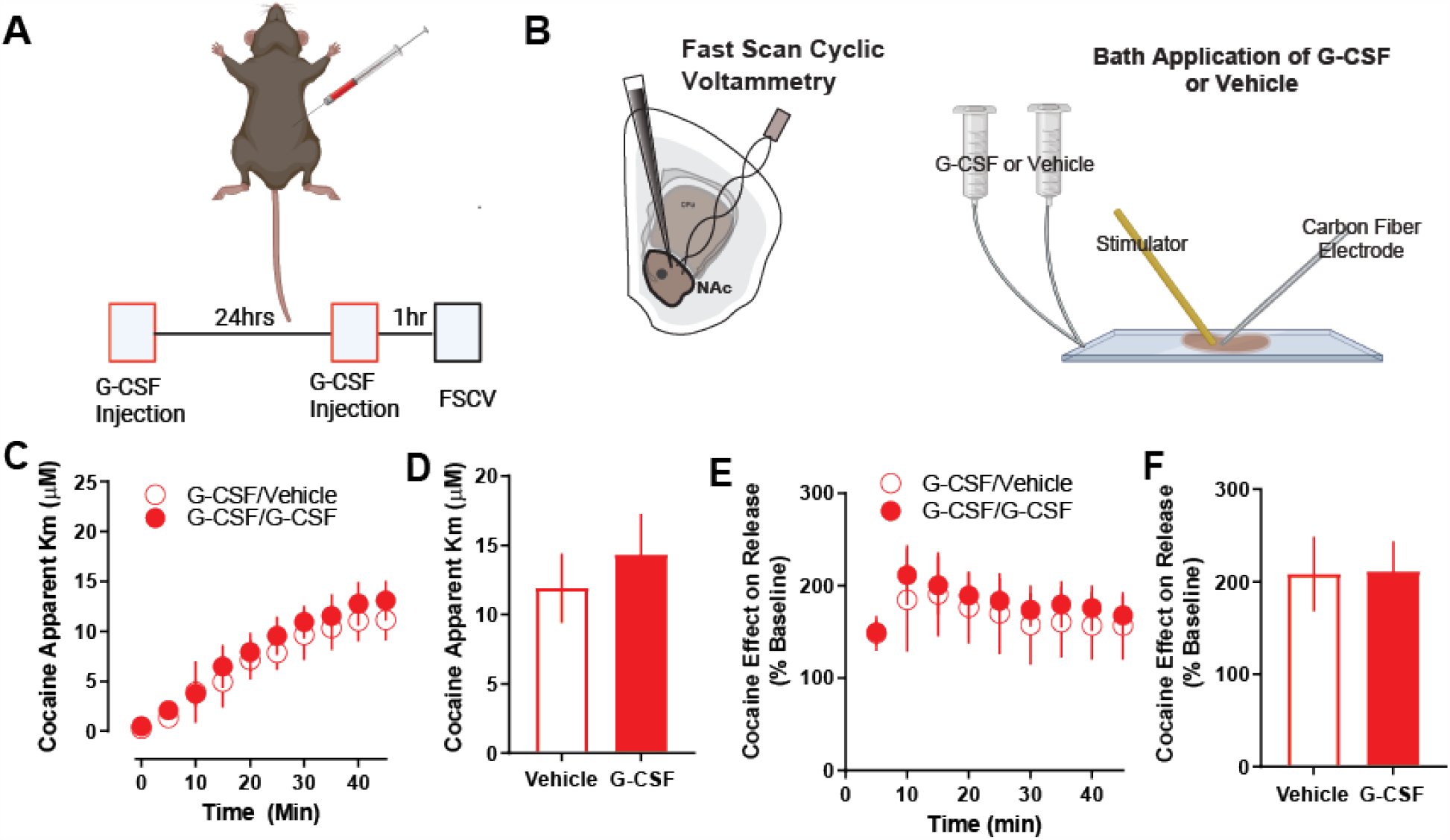
No effect of G-CSF bath application on cocaine in G-CSF pre-treated animals. **A**. Male C57BL6/J mice were injected with G-CSF (50μg/kg, IP) twice – 24 hours before recording and again one hour before recording. **B**. Fast scan cyclic voltammetry was run, and G-CSF or vehicle was applied via bath application to the NAc slices for 1 hour prior to cocaine application (n = 3 per group). **C**. G-CSF bath application had no effect on cocaine effects at DAT as measured by apparent Km. **D**. Group averages also showed no difference in apparent Km measures. **E**. Cocaine effect on dopamine release as presented as a percent of baseline dopamine release. **F**. G-CSF bath application also had no effect on cocaine effects on release following G-CSF pre-treatment. Data represented as mean +/- S.E.M.

Together these data suggest that G-CSF enhances cocaine effects in the NAc through a mechanism that is specific to release. Further, this effect is directly in the NAc as it can be induced via direct G-CSF application to NAc slices, an effect that is occluded via prior G-CSF injection - suggesting that the mechanism for systemic and local G-CSF application is the same.

## Discussion

Here we show that G-CSF alters the pharmacodynamic properties of cocaine in the NAc. We previously found that prolonged treatment with cocaine leads to significant increases in levels of circulating G-CSF as well as increases in expression of G-CSF and its receptor in the NAc of mice^5,6^. Additionally, injections of G-CSF enhanced self-administration and conditioned place preference for cocaine; however, the neural mechanism for these effects was unclear^6^. Here we show that G-CSF alters cocaine effects at dopamine terminals, where systemic injections or direct application enhances dopamine release in the presence of cocaine. This effect occurs through a release-based mechanism, rather than through effects on the dopamine transporter. Importantly, this release-based mechanism could not be explained purely by changes in evoked release at baseline and represented additive enhancement of cocaine effects on release. The effect occurs via direct actions of G-CSF in the NAc as it could be recapitulated by acute bath application of G-CSF to dopamine terminals. Together, this work adds important mechanistic insight into the role of G-CSF in cocaine use disorder and defines how enhanced G-CSF levels following cocaine exposure could be contributing to its development.

In recent years, there has been a growing understanding that interactions of the immune system and the central nervous system are likely playing a causal role in the pathophysiology of multiple psychiatric illnesses. As a result of observational clinical studies and translational animal studies, we are moving closer to understanding the mechanistic links between immune dysregulation and psychiatric pathology. While the role of neuroimmune interactions is more extensively studied in other psychiatric conditions, such as depression, schizophrenia, and autism^25–27^, recent work has identified neuroimmune mechanisms as potential critical mediators of brain and behavior in substance use disorders^2,3^. While the mechanisms underlying neuroimmune signaling are complex^28^, signaling through soluble cytokines is the best studied and most probable mechanism for driving neuroimmune effects in neuropsychiatric disorders^2,29^. In the striatum signaling through inflammatory cytokines alters homeostatic synaptic plasticity^7,30,31^. Early research into these cytokine dopamine interactions has focused on pro-inflammatory cytokines (e.g. Interferon-α^32,33^, IL-6^34,35^, TNF-α^36,37^, etc.) and suggest that they represent a potentially important means of fine-tuning dopamine signaling to drive appropriate responses to rewarding and salient stimuli. Accordingly, there is a wide body of literature demonstrating that cytokines have effects on dopamine signaling in the brain and can alter transcription within dopamine neurons, dopamine synthesis, and dopamine release and reuptake in behaviorally important ways^38–41^ – indeed, these changes in dopamine dynamics are linked to the behavioral effects of cocaine^17,20,42^.

While there is evidence that other proinflammatory cytokines exert their effects through altering dopaminergic function, much of the previous work on this area has suggested that these factors reduce, rather than increase, dopamine release and regulation. While G-CSF has been shown to have effects on neurons throughout the brain, it seems to exert particularly robust effects on the dopamine system, where it regulates dopamine neuron development, cell health, and dopamine release. For example, treatment with G-CSF is neuroprotective of dopamine neurons in both basic^43,44^ and clinical^45^ studies of Parkinson’s disease, and G-CSF enhances activation of immediate early genes in dopaminergic neurons^46^. Thus, the differences between other inflammatory cytokines and G-CSF on dopamine neurons could be due to 1) G-CSF exerting different effects on these neurons through distinct signaling cascades or 2) differences in the way that dopaminergic function was assessed in the prior studies. For example, cytokines may be acting to refine signaling and enhance signal to noise, which would result in reductions in measurements that assess tonic dopamine levels, but enhancements in dopaminergic measurements that assess evoked release that mimics responses to salient stimuli *in vivo* – such as voltametric measures that were used here to assess sub second dopamine release following an electrical stimulation. Moving forward it will be important to assess these complexities in the effects of cytokine signaling on dopaminergic regulation.

Further, identifying where in the brain immune signaling molecules are acting to exert their effects on cocaine action is important, given that many of these cytokines act broadly throughout the body and brain. Here we show that the effect of G-CSF on cocaine occurs directly within the NAc. By bath applying G-CSF to coronal slices containing the NAc we show here that cocaine’s effects on dopamine release are altered, even though the terminals were dissociated from their cell bodies. While it is clear that these effects originate in the NAc, it is unclear whether they are occurring through direct actions on dopamine terminals, or through another microcircuit-based mechanism. Dopamine release from terminals in the striatum is regulated by a wide range of neurotransmitter systems including acetylcholine, glutamate, GABA, various peptide systems and other types of signaling molecules (for review see Nolan et al., 2020^47^). While we hypothesize that these effects do indeed occur through direct terminal regulation, glutamatergic neurons in the mPFC that project to the NAc are also known to play a role in the effects of G-CSF^6,10^. Thus, it is also possible that G-CSF is acting on cortical terminals in the NAc which alters NAc activity and dopamine release through a microcircuit.

Another important consideration when assessing the effects of G-CSF in the brain is biological sex. While this study focused on male subjects, G-CSF mediated enhancement of dopamine release has been observed in females; however, in females this originates from a different mechanistic basis^5^. While injections of G-CSF enhanced cocaine conditioned place preference and increased cocaine’s ability to increase dopamine levels in both male and female mice, in females G-CSF treatment enhanced cocaine effects through altering cocaine interactions with the dopamine transporter - with no effects on release^5^. This is in stark contrast to the data from males presented here that shows that G-CSF effects are through a release-based mechanism and do not alter transporter-mediated clearance. This is consistent with data across fields showing that, in some cases, even when there are similar behavioral phenotypes between males and females that they can be driven by different molecular mechanisms^48–50^.

Regarding translational potential, targeting of neuroimmune interactions is an area of growing interest in the neurobiology of substance use disorders, and there is considerable promise for targeting these systems to modulate the symptoms of addiction. G-CSF signaling enhances cocaine taking and seeking^6^, increases dopamine release in the NAc^4,5^, alters learning for rewarding stimuli^4^, and significantly alters protein expression in multiple limbic structures including within dopaminergic neurons^10,41^. Thus, G-CSF signaling seems to be a potent regulator of dopamine neurons that allows for regulation of dopamine release and behavioral effects, without direct agonist/antagonist effects on dopamine release in the absence of other stimuli. Importantly, we also have demonstrated that treatment with G-CSF during abstinence after cocaine self-administration can reduce subsequent drug seeking behavior^10^.

Given the high value of neuroimmune interactions in developing new therapeutic strategies for substance use disorders and other psychiatric conditions, and the clear evidence for G-CSF in altering key limbic structures and behaviors, defining its precise signaling mechanisms and effects within midbrain dopamine neurons is critical. Further, G-CSF specifically is a promising target as compounds that modulate its levels or function are already FDA approved for use in clinical populations. However, to translate findings such as these into individuals with substance use disorder, it is imperative to understand the complex factors that influence its actions at a molecular and level.

## Acknowledgements

Cocaine hydrochloride was provided by the NIDA Drug Supply Program. Diagrams in all figures were created with Biorender.com with full permission to publish.

## Author Contributions

LJB and KRE collected data, analyzed data, made figures, and wrote the manuscript. KEL, AO, analyzed data, made figures, and wrote the manuscript. ESC and DDK conceptualized the manuscript, analyzed data, made figures and wrote the manuscript.

